# Photothermal Dye-based Subcellular-sized Heat Spot Enabling the Modulation of Local Cellular Activities

**DOI:** 10.1101/2021.12.19.471878

**Authors:** Ferdinandus, Madoka Suzuki, Yoshie Harada, Satya Ranjan Sarker, Shin’ichi Ishiwata, Tetsuya Kitaguchi, Satoshi Arai

## Abstract

Thermal engineering at microscale such as the control and measurement of temperature is a key technology in basic biological research and biomaterials development, which remains challenge yet. Here, we engineered the polymeric nanoparticle, in which a fluorescent temperature sensory dye and a photothermal dye were embedded in its polymer matrices, termed **nanoHT**. When a near infrared laser at 808 nm is illuminated to the particle, it enables to create the subcellular-sized heat spot in a live cell, where fluorescence thermometry allows the read out of the temperature increment concurrently at individual heat spots. Owing to the controlled local heating, we found that the cell death of HeLa cells was induced at the certain temperature at rate of a few seconds. It should be also noted that the cell death was triggered from the very local heat spot at subcellular level. Furthermore, **nanoHT** was applied for the induction of muscle contraction of the C2C12 myotube by heat. We successfully showed that the heat-induced contraction took place at the limited area of a single myotube according to the alteration of protein-protein interactions related to the contraction event. These studies demonstrated that even a single heat spot provided by a photothermal material could be very effective in altering cellular functions, paving the way for novel photothermal therapies.

## Introduction

Heating technology at nano/microscale has been indispensable in various research fields such as material engineering and biological sciences^1,2,3^. More specifically in biological research, it should be noted that inside of a single cell is filled with a huge variety of temperature-sensitive elements such as chemical reactions, fluidity of cellular membranes, and flexible structure of biomacromolecules. Therefore, further advances in heating methods are expected because of its great potential to contribute the analysis of thermodynamic elements in biology and accelerate the development of effective biomaterials^4^. To date, magnetic- and optical heating for biospecimens has been established as means applicable at cellular and tissue levels^5,6^. Yet, the optical one is superior to the other from the viewpoint of spatiotemporal resolution^1^. For instance, an optical microscopic system with a near infrared laser at 1460 nm was fabricated to generate the temperature increment at the localized spot through activating the vibration of water molecules, termed IR-LEGO system^7^. However, the spatial resolution of the temperature distribution provided with IR-LEGO is still limited to a size of the laser spot^7^. To achieve the heat spot with more narrow spatial resolution, a nano-sized photothermal agent is required, that can absorb the light energy and convert into heat. So far, intensive efforts are dedicated to develop inorganic materials^8^, nanocarbons^9^, semi-conductive polymers^10^ and organic dyes^11^, termed a nanoheater. While under microscopic observation, the laser irradiation with an appropriate wavelength to a nanoheater could produce the heat spot at the tiny size. As another important aspect for precisely controlled heating, the temperature sensing in the proximity to nanoheater, possibly at zero-distance from the heat spot, is essential due to the short-lived heat propagation^6,12^. Recent years, nanomaterials that possess both of heating and thermometry functions were developed as a nanoheater-thermometer system^13,14,15^. Some of them were further demonstrated in photothermal therapy with the *in vivo* thermometry including animal studies^16^. However, despite numerous studies including the simulation at nanoscale, it has been scarcely argued how a single dot of a nanoheater generates the temperature increment and its distribution under the environment of a live cell^17,18,19^. Consequently, how a single nanoheater alters cellular activities in real-time has not been also figured out at single cellular level. Other than that, a question here is whether an even single nanoheater is efficient to affect cellular functions at subcellular level.

Here, we engineered a polymeric nanoparticle with the ability of heat generation and temperature sensing in individual heat spots. More specifically, a fluorescent temperature sensory dye was embedded as well as a photothermal dye into a polymeric nanoparticle where the temperature increment at the heat spot could be captured with the thermometry. Successes of several fluorescent thermometers of late have been reported that are capable of reporting the intracellular temperature as detectable fluorescence signals^20^. While the accuracy of fluorescence thermometry remains challenging, it still benefits the combination with several fluorescent indicators to image cellular events with concurrent thermometry. Using our heating technology, we examined how a single heat spot induced the heat-triggered cell death of HeLa cells from viewpoints such as temperature threshold and the alteration of intracellular Ca^2+^ and ATP dynamics. We also attempted to manipulate the muscle contraction in C2C12 myotube by the subcellular local heating. Through several imaging studies, we show how the tiny heat spot behaves with the alteration of cellular functions.

## Results

### *In vitro* Characterization of nanoHT

We designed a polymeric nanoparticle containing dyes for sensing temperature and generating heat via photothermal effect, termed the nanoheater-thermometer (**nanoHT**) (Fig. 1A). To preserve more accurate temperature sensing, both of temperature sensitive and -less sensitive dyes, europium tris-(dinaphthoylmethane) complex (EuDT) and courmarin102 (C102) respectively, were embedded into the polymeric particle^21^. Both of dyes can be excited at a blue laser (405 nm) simultaneously and its fluorescence emission recorded separately (C102: 430-455 nm and EuDT: 575-675 nm), allowing fast ratiometric temperature monitoring (Fig. 1B). As a photothermal agent, a near-infrared (NIR) absorbing dye was chosen because it is compatible to be incorporated into a hydrophobic polymer matrix. Common NIR absorbing dyes such as linear cyanine derivatives (IR780 and indocyanine green) are widely used in the biomaterial development but encounters difficulties in its photostability^22^. On the other hand, the photostability of phthalocyanines is predominant over that of linear ones. In addition, metallo-phthalocyanines with heavy metals prefer the non-radiative excited state relaxation after the absorption of a NIR laser light, leading to the efficient photothermal effect^23^. Considering the appropriate wavelength not to interfere with the fluorescence emissions of C102 and EuDT, a vanadyl tetra t-butyl naphthalocyanine (V-Nc) was finalized which can be illuminated by a common NIR laser at 808 nm^24^. The poly(methyl methacrylate-*co*-methacrylic acid)(PMMA-MA) based polymeric nanoparticle was prepared via the nanoprecipitation method, in which three dyes of C102, EuDT and V-Nc were incorporated to the matrix^25^. The diameter of resulting particles was measured to be 153±51 nm by dynamic light scattering (Fig. 1C and Supplementary Fig. S1). We further characterized the ability to sense temperature and release heat in the cuvette. As expected, the fluorescence intensity of EuDT declines as temperature increases while that of C102 was scarce (Fig. 1D). The calibration curve on the fluorescence intensity vs. temperature correlation was also obtained showing the linear slope between 35 to 45 °C at least. Regarding the ability of the heat release, the rise in temperature in a suspension of **nanoHT** in the cuvette upon a NIR laser (808 nm, CW) irradiation was examined using a thermocouple while the temperature increment was negligible in water as a control experiment (Fig. 1E). The photothermal conversion efficiency of **nanoHT** was estimated to be 35% according to the previous literature^26^. In the end, the property to generate reactive oxygen species (ROS) was evaluated using a toolkit sensing non-specific ROS (H2DCFDA) (Supplementary Fig. S2). This is because our study is aimed at investigating how the only heat generated by **nanoHT** contributes to cellular functions and thereby the effects by the other elements like ROS should be avoided as much as possible. As compared with the gold-nanorod (AuNR) as a representative photothermal material, **nanoHT** indicated the quite low ability of ROS production, which would be a great advantage through this study.

**Figure 1.**
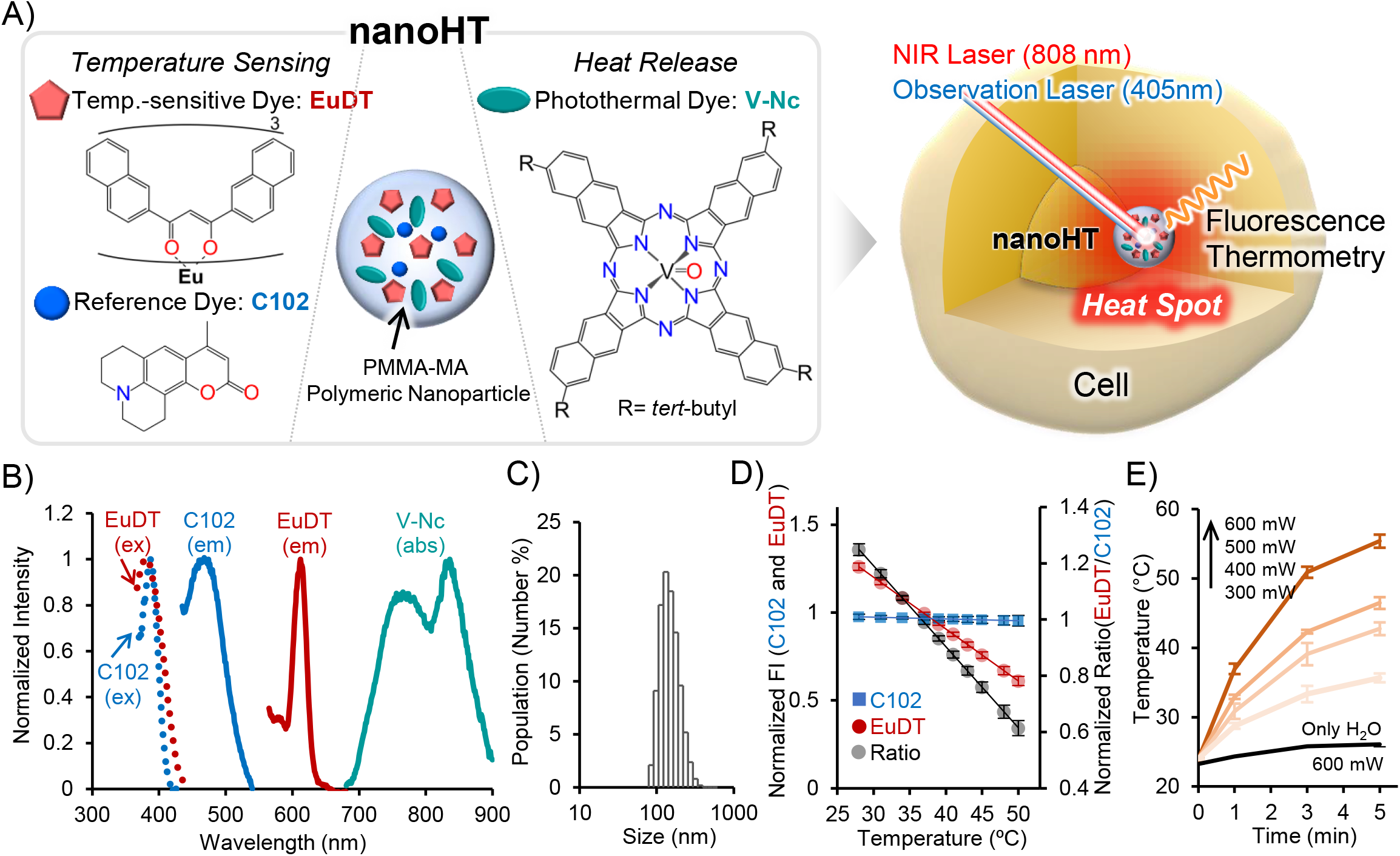
Characterization of **nanoHT** with the ability of heat release and temperature sensing. A) Schematic illustrations of **nanoHT** and its controlled heating inside a cell. B) Excitation and fluorescence spectra of C102 and EuDT, and absorption spectrum of V-Nc in **nanoHT**. C) DLS measurement of **nanoHT**. The average of diameter: 153 ± 51 nm (mean ± SD). D) The normalized fluorescence intensity (FI) values of C102 and EuDT were plotted against temperature as the 1^st^ axis, and the ratio value (EuDT/C102), which is normalized to that of 37 °C, as the 2^nd^ axis. Error bars, SD (n=3). E) The evaluation of the heating ability of **nanoHT** suspension in the cuvette by the irradiation of a NIR laser (808 nm). Error bars, SD (n=3).

### Validation of nanoHT under microscopic system

We investigated the feature of **nanoHT** under microscopic observation. The diluted solution of **nanoHT** was casted on the glass so that only a few **nanoHT**s could be observed at a microscopic observation area. The fluorescence was recorded in two channels for C102 and EuDT at the same time (Fig. 2A). Firstly, we validated the temperature sensing ability of **nanoHT** through heating up surrounding medium. To warm up the surrounding medium, we adopted the technique in which a NIR laser (980 nm) coupled with an iron agglomerate enables to create reversible temperature gradient quickly^27^. For reference, V-Nc embedded as a photothermal dye absorbs a light at 980 nm scarcely though the water molecule can absorb it and contribute the temperature increment of the medium partially. When the shutter of a 980 nm laser being turned on, the fluorescence intensity of EuDT declined in response to the temperature increment while that of C102 was scarce. And then, when being off, it returned to the basal level, showing a step-like change of fluorescence (Fig. 2B). The depth of the step differs depending on the distance between the heat spot and **nanoHT** (ROI1, 2 and 3). While the temperature in the medium was varied by the temperature controller of the microscopic chamber, the calibration curve between temperature and the normalized ratio (EuDT/C102) was obtained as shown in Fig. 2C. Hereafter, the normalized ratio value was defined where the ratio value of EuDT to C102 was divided by that at 37 °C. The temperature sensitivity and its resolution (2^nd^ axis) are estimated to 1.85 %/°C and 0.3-0.8 °C, respectively, which are compatible with previous polymeric nanoparticle type-fluorescent sensors^20^. Using this calibration curve, the difference of the normalized ratio could be converted to the temperature increment (Δ*T*) (Fig. 2D).

**Figure 2.**
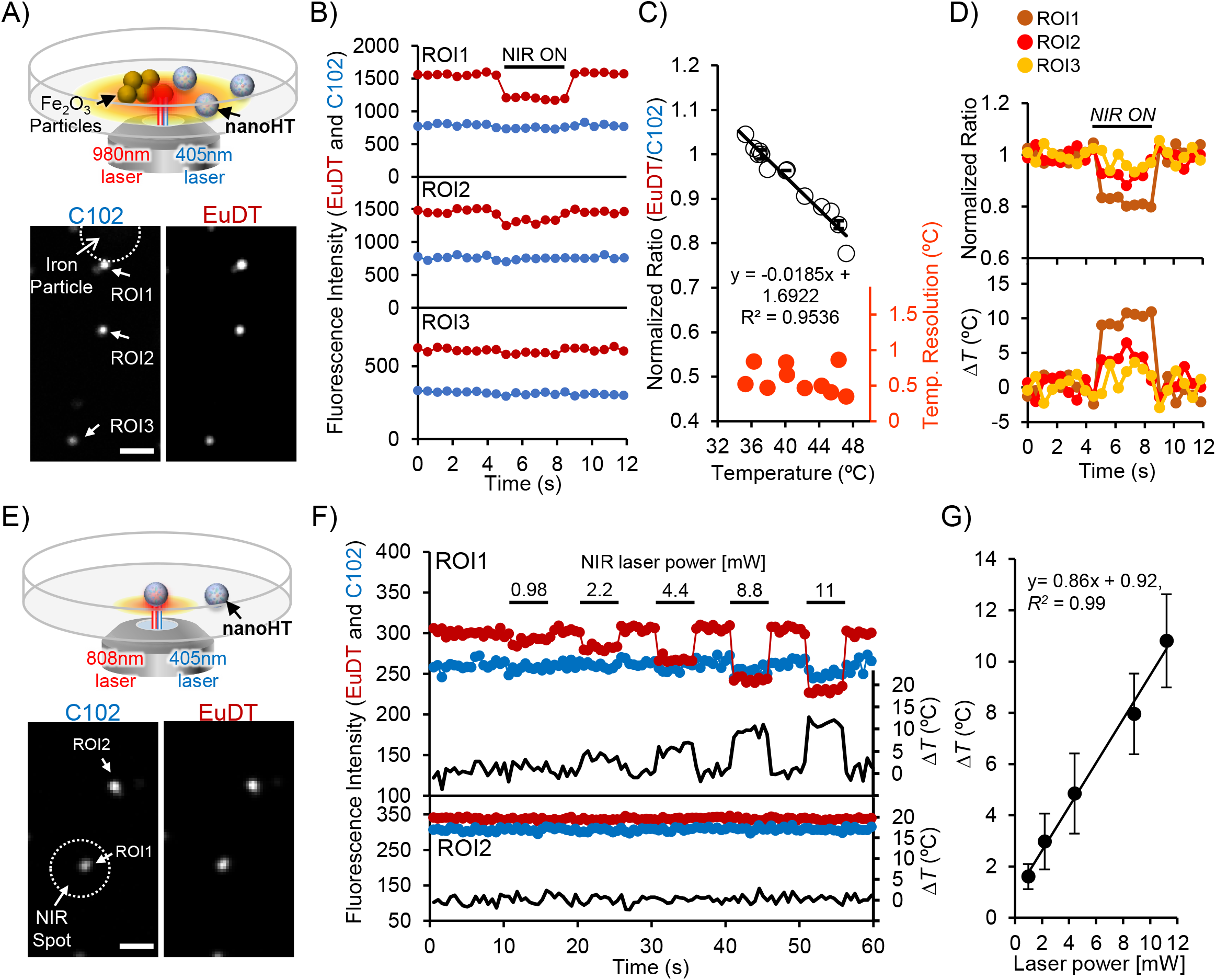
Validation of **nanoHT** with the ability of heat generating. A) Schematic image of a means to validate the temperature sensing ability of **nanoHT** under the microscope with a NIR infrared laser (980 nm). Scale bar: 5 μm. B) The mean fluorescence intensities of C102 and EuDT at each ROI as shown in A) were plotted every 0.56 sec in the time course (5 sec NIR laser stimulation). C) The calibration curve of **nanoHT** against temperature obtained under the microscopic observation. Error bars, SD (n=3). D) The normalized ratio (EuDT/C102) was converted to the temperature increment profile (Δ*T*) using the calibration curve. E,F) Validation of the heat generating ability of **nanoHT** using an 808 nm laser. A NIR laser was only illuminated to a white circle spot. The mean fluorescence intensity at each ROI was plotted in the time course while a NIR laser stimulation is performed during 5 sec at different laser powers (0.98-11 mW). G) The average of temperature increment provided by **nanoHT** (error bars, SD. n = 10) was plotted at each 808 nm laser power. Solid line showed the linear fit.

Switching from a 980 to 808 nm NIR laser, we further examined the heating ability of **nanoHT** by the irradiation (Fig. 2E). An 808 nm laser scarcely warms up the medium and is suitable for the absorption of V-Nc as a photothermal agent. When an 808 nm laser was illuminated to a single dot of **nanoHT** and the on/off of the shutter repeated, the fluorescence of **nanoHT** exhibited a step-like behavior, which is similar to Fig. 2B (Fig. 2F). Regarding the temporal resolution, the heat spot could be generated and erased within less than one frame shot during timelapse experiments (images were taken every 0.56 sec). Varying a laser power created the step with different depths, generating the different temperature increment at the heat spot (Fig. 2F). Importantly, it was found that **nanoHT** located outside the NIR laser spot (ROI2) rarely exhibited the temperature increment (lower panel in Fig. 2F). What we stress here is that the combination of **nanoHT** and an 808 nm laser enables the targeted and fast heating on the spot. The average of Δ*T* provided with **nanoHT** was plotted against different laser powers (Fig. 2G). The variation of its increment at the same laser power was bigger than that of the accuracy of thermometry effectively, suggesting its variation would be due to the size variation (153 ± 51 nm) of **nanoHT**.

### Investigation of the temperature distribution by nanoHT

When **nanoHT** applied for HeLa cells, it was uptaken into the cell through the endocytic pathway without the significant cell toxicity (Supplementary Fig. S3). The co-localization test with organelle trackers suggests that **nanoHT** was localized to acidic organelles such as endosomes and lysosomes (Supplementary Fig. S4). In the same way as the glass plate (Fig. 2E-G), an 808 nm laser was irradiated to a single **nanoHT** in a live HeLa cell during microscopic experiments. Likewise, step-like responses of fluorescence of **nanoHT** and its laser power dependency of depths could be observed (Fig. 3A-C). Moreover, varying the on/off timing of the shutter of an 808 nm laser, the different temporal pattern of the temperature increment could be generated as shown in the Fig. 3D (every 2, 5, and 20 sec). The loss of the heating ability upon the repeating stimulation was negligible under these conditions at least. Notably, the escape of **nanoHT** from acidic organelles was observed during a couple of seconds heating, supported by the results that the fluorescence derived from acidic organelles tracker was diminished after heating (Fig. 3E). Presumably, heat induced the collapse of the endosomal membrane according to literatures regarding several types of photothermal nanomaterials^28^.

**Figure 3.**
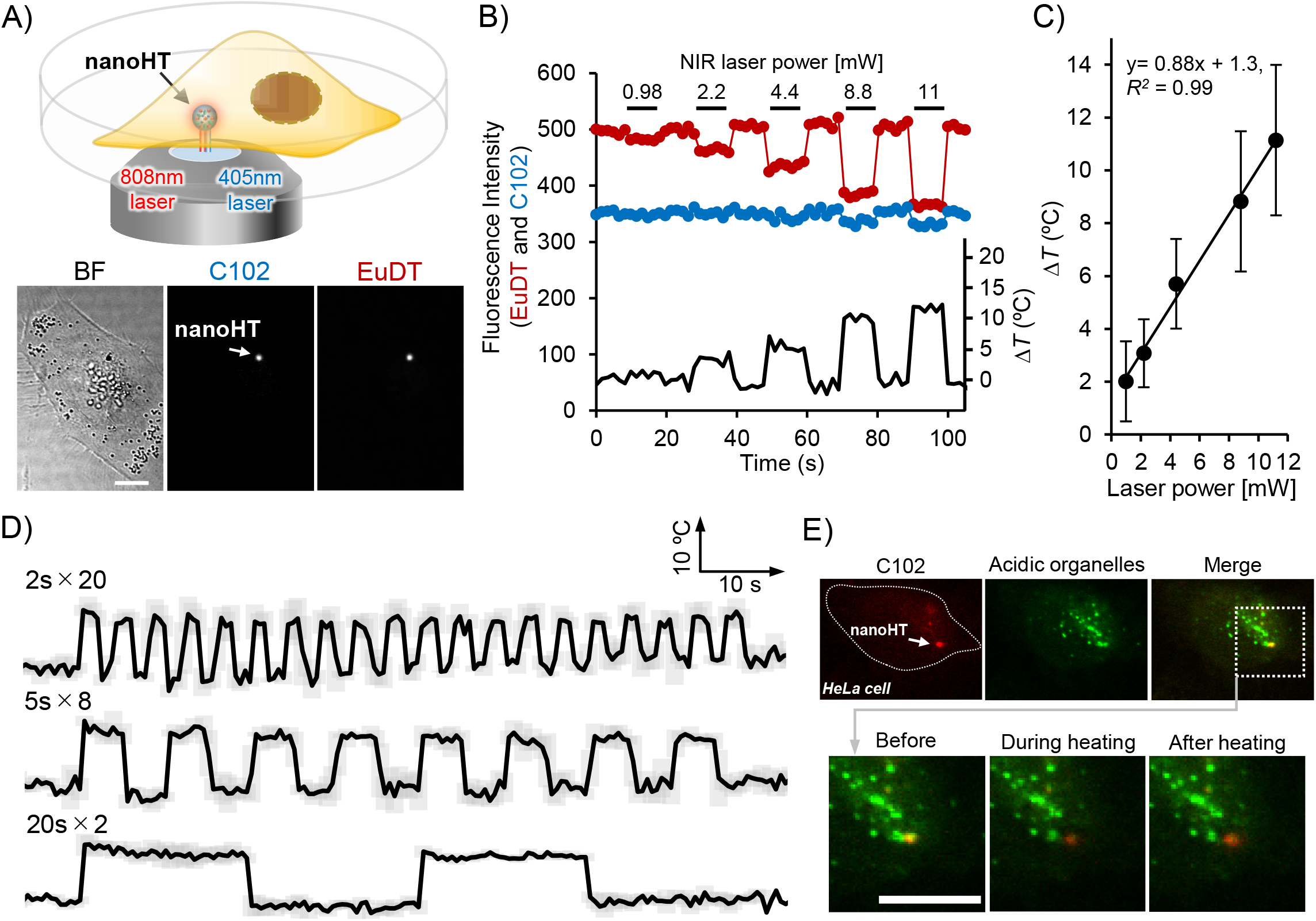
Validation of **nanoHT** in HeLa cells. A,B) Validation of the heat generating ability of **nanoHT** using an 808 nm laser. The mean fluorescence intensity of **nanoHT** in A) was plotted in the time course while a NIR laser stimulation was performed during 5 sec at different laser powers (0.98-11 mW). Scale bar: 10 μm. C) The average of temperature increment provided by **nanoHT** (Error bars, SD. n = 12) were plotted at varying 808 nm laser powers. Solid line showed the linear fit. D) The different temporal pattern of the temperature increment created by **nanoHT**. The mean of normalized ratio of **nanoHT** with SD (n=3) was plotted in the time course. E) Colocalization test with a lysosome tracker in the upper panel (Red: C102, Green: lysosome tracker to stain acidic organelles). Enlarged view of the region surrounded by a dashed square before, during and after heating. Scale bar: 10 μm.

We next addressed the spatial distribution of temperature provided by **nanoHT**. For the analysis of the temperature distribution, the other fluorescence temperature sensor, a blue fluorescent protein (BFP) was added to the medium together with **nanoHT**, that can map out the temperature in the surrounding medium (left panel: Fig. 4A, Supplementary Fig. S5)^29^. As a merit using BFP, it is also applicable for the temperature mapping in cytoplasm of a live cell through the gene expression of the same sensor (right panel: Fig. 4A). Firstly, the movement of **nanoHT** was examined in the glass dish and inside a live cell with varying the laser power using the particle tracking software. As it turned out, the moving distance of **nanoHT** in a live cell was much longer than that in the dish within the same duration (Fig. 4A). It could be reasoned that **nanoHT** moves relatively freely inside the cell while **nanoHT** sticks to the glass bottom. The further analysis of the velocity of **nanoHT** (μm/s) also supported that **nanoHT** in the cell is faster than that in the glass dish. As the other aspect, the velocity of **nanoHT** inside the cell even at the maximum temperature increment is smaller than that of the conventional nanoparticles transported by the motor protein (0.32 μm/sec at 36°C)^30^. In addition, Oyama et al. described previously that the velocity of the nanoparticle during the transportation via motor proteins exhibited temperature-dependent manner^30^. As such, the temperature dependency of the velocity of **nanoHT** was evaluated but the significant trend could not be observed (Fig. 4B). Considering these results, almost **nanoHT** escaped from acidic organelles after heating, followed by the floating inside the cytoplasm and governed by the Brownian motion. In such case, capturing the exact position of **nanoHT** would be considerably difficult due to fast random walk at this time scale, and thus its temperature dependency was likely to be unclear. There might be another possibility that some of them stick to the intracellular components inside cell non-specifically and consequently difficult to move^31,32^.

**Figure 4.**
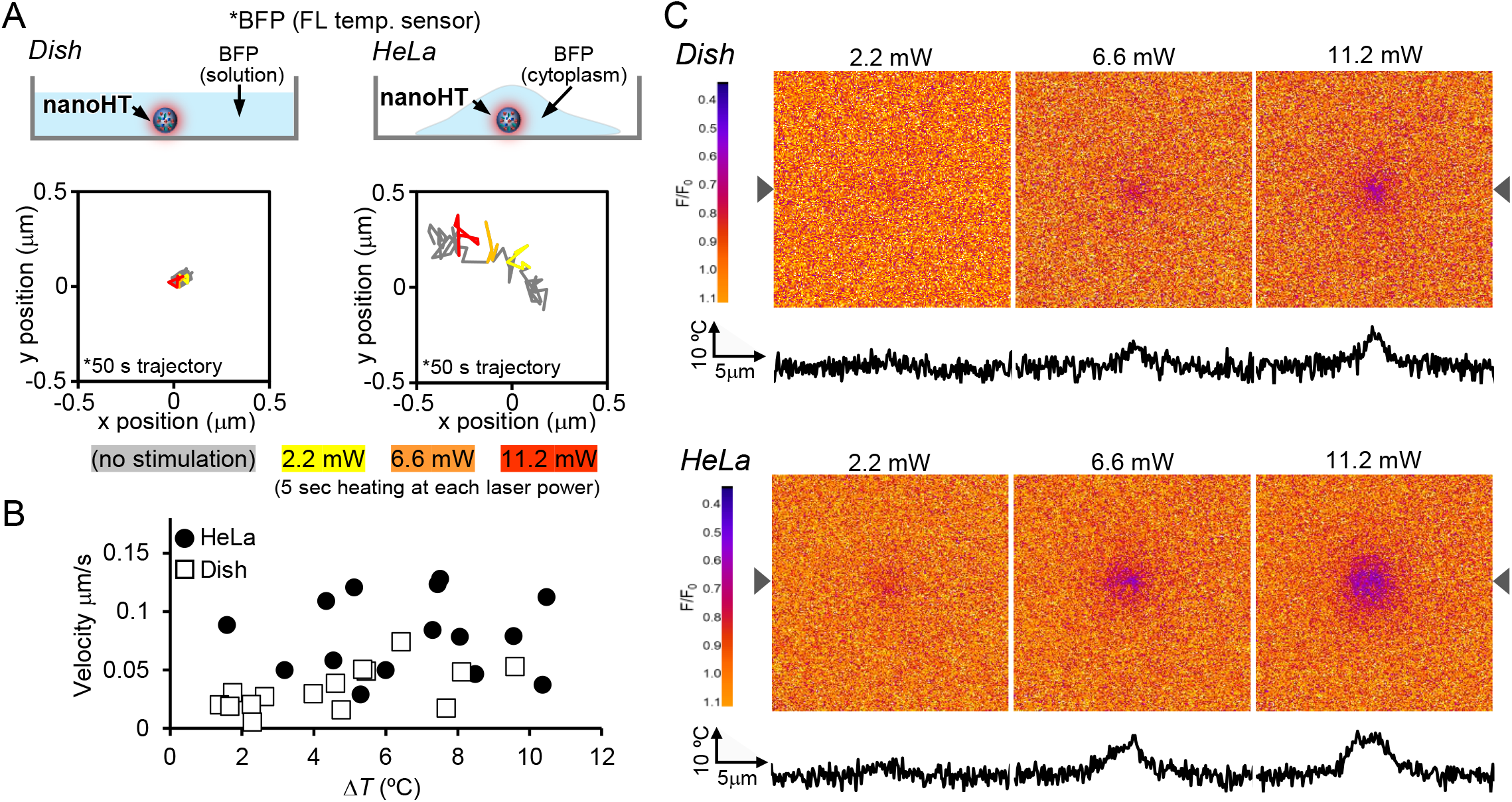
Evaluation of temperature distribution provided by **nanoHT** in a HeLa cell and in the dish. A) **nanoHT** was located at the surface of the dish filled with the blue fluorescent protein (BFP) solution while **nanoHT** was taken into the HeLa cell expressing BFP. The trajectory of **nanoHT** was depicted at lower panel in the dish and HeLa cell, respectively. During the 50 sec tracking, the NIR laser stimulation was performed at three different powers (2.2, 6.6, and 11.2 mW) during 5 sec. B) The velocity of **nanoHT** (μm/s) during heating was plotted at different temperature in the dish and HeLa cell. C) The analysis of temperature distribution generated by **nanoHT** using BFP at different laser powers (2.2, 6.6 and 11.2 mW). The grouped stacked images during 5 sec heating were divided by the image before heating. The triangle marks indicated the position of the line profile as shown at the bottom of each image.

In the end, the analysis using a BFP temperature sensor proved the different spatial pattern of temperature distribution in the dish and inside the cytoplasm (Fig. 4C). As expected, the bigger the temperature increment was, the larger the spatial distribution was in both cases. As the distinguishable point, the spatial distribution of temperature in the cytoplasm was larger than that in the dish apparently. These temperature mapping images were obtained from the group-stacked analysis during 5 sec heating and represented the accumulated history of the temperature change occurring over a period of 5 sec. In other words, it could be ascertained that the discrepancy of the distribution should reflect on the ease of the movement of **nanoHT** in different environments. Inside a cell, **nanoHT** is possible to offer the thermal effect at subcellular scale of a couple of microns during a few seconds.

### Rapid induction of the cell death in HeLa cells

By local heating with **nanoHT**, we tested how the heat induced the cell death of HeLa cells. To identify whether the heat-induced cell death occurs or not, HeLa cells were stained together with Apopxin Green to detect the PS (phosphatidylserine) and a membrane-impermeable propidium iodide (PI) dye to stain nucleus. The former is frequently used for apoptosis detection because the PS is transferred to the outer leaflet of the plasma membrane while the latter for detection of necrosis or apoptosis at late stage involved in the rupture of plasma membrane. With varying the different laser power (from 8 to 11 mW), we found the enhancement of the fluorescence of Apopxin Green within a few seconds at a certain temperature increment (Fig. 5A and B). More interestingly, the apoptosis marker (PS marker) appeared to gather nearby the local heat spot. In addition, cells with the enhancement of Apopxin Green by heating also showed the increase of the fluorescence of PI and the bleb formation after 10 min (Fig. 5C). It is assumed to satisfy requirements partially for the identification of apoptotic-like cell death (the detail will be argued in the discussion part). The correlation between Δ*T* of **nanoHT** and the normalized fluorescence of Apopxin Green clarified the threshold temperature of the cell death, which was estimated to be around 11.4 °C (=Δ*T*, base temperature: 37 °C) (Fig. 5D). The dynamics of calcium ion (Ca^2+^) and the Apopxin Green were further imaged at the same time and at the same cell (Fig. 5E). Once the heat generated by **nanoHT**, the elevation of an intracellular Ca^2+^ level was induced from the heat spot at the early stage. The possible explanation of the local Ca^2+^ elevation is that local heating at subcellular level perturbed functions of mitochondria and endoplasmic reticulum (ER) as an intracellular pool of Ca^2+ 33^. Particularly, previous studies hypothesized that heat stress caused the perturbation of electron transport chain (ETC) of mitochondria and the increase of mitochondrial membrane permeability, resulting in the cell death with the leak of the Ca^2+ 34^. Notably, the elevation of ROS was also observed during heating (Supplementary Fig. S6). Since **nanoHT** was identified to be an ineffective photosensitizer for ROS production (Supplementary Fig. S2), ROS should not be provided directly from **nanoHT** but derived from the effect by the photothermal perturbation to mitochondria. To the best of our knowledge, it is the first case to capture the cell death triggered by the sub-cellular sized heat spot in real-time with concurrent thermometry. Also, it is worth noting that the induction of the cell death at the rate of a few seconds appears to be a rare case compared to previous cases that took a few hours^35^.

**Figure 5.**
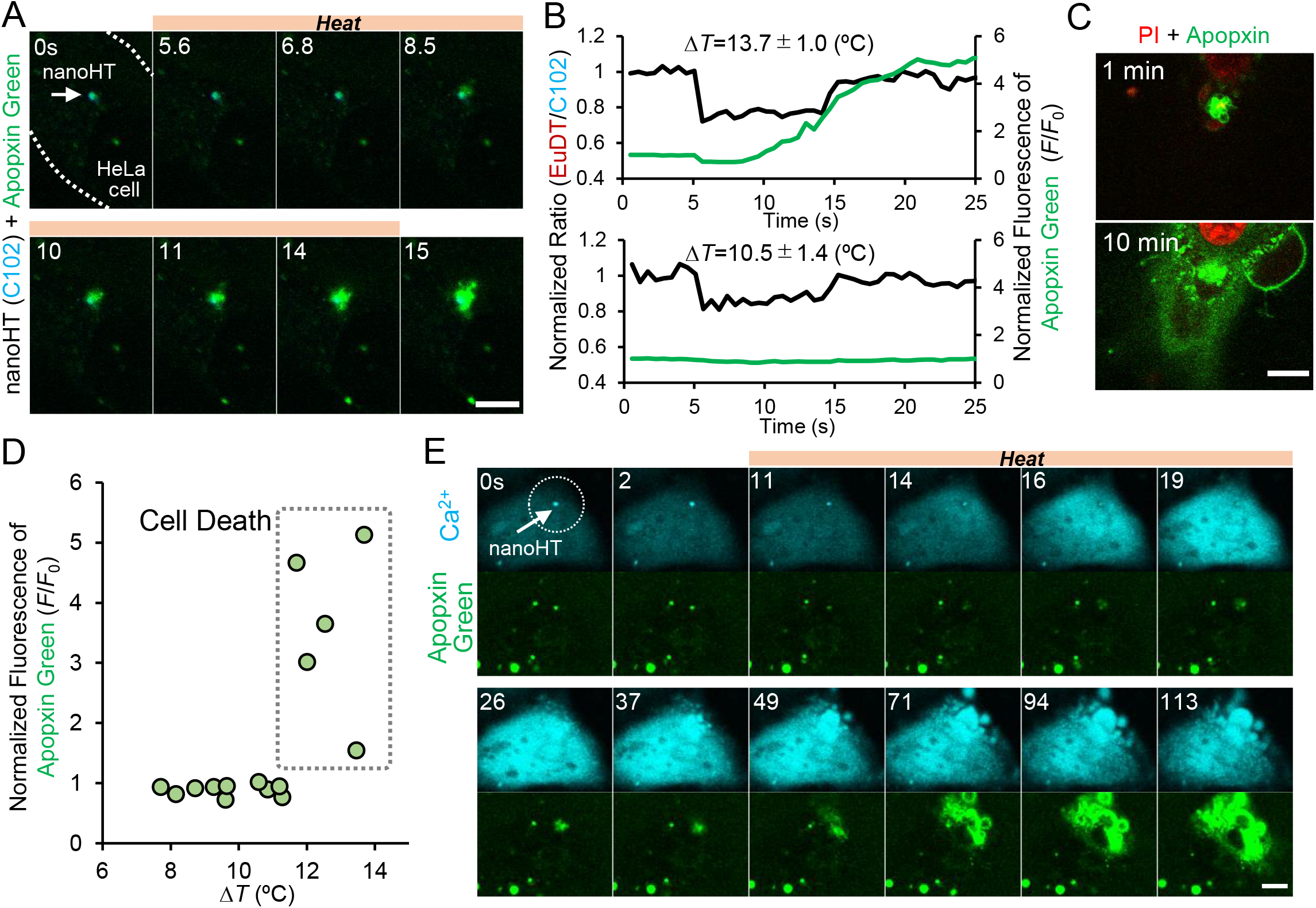
Heat-triggered cell death by **nanoHT**. A) Dual imaging of Apopxin green (apoptosis marker) and **nanoHT** (Blue: C102) in a HeLa cell. B) The time course of the normalized ratio of **nanoHT** and fluorescence of Apopxin green nearby the heat spot (a NIR stimulation during 10 sec). The temperature increments were estimated by the calibration curve. C) Images of HeLa stained with Apopxin green and PI (for detection of necrosis or late stage of apoptosis) after heating (1 and 10 min). D) The correlation between temperature increment of **nanoHT** and the enhancement of Apopxin green (*F*/*F*_0_). Laser power was varied from 8.8 to 11.2 mW. E) Dual imaging of Ca^2+^ (B-GECO) and Apopxin green in a HeLa cell. Elapsed time is shown to the top left of each image in A) and E). Scale bars: 10 μm.

We next sought to elucidate the heat-induced impact on cells more from the viewpoint of the dynamics of adenosine triphosphate (ATP) as a key in energy metabolism. To image intracellular ATP, MaLionG and mitoMaLionR as genetically encoded fluorescent ATP sensors were expressed to HeLa cells for monitoring cytoplasmic and mitochondrial ATP, respectively^36^. As a result, under almost cases to induce the cell death, mitochondria were broken down into a piece remarkably while mitochondrial ATP level was declined (Fig. 6A and Supplementary Fig. S7). The fragmentation of mitochondria in the conjunction with the irreversible ATP depletion was characterized in apoptotic-like cell death^37^. Subsequently, we investigated how ATP dynamics was altered by heat stress below the threshold temperature for the cell death. When local heating applied for 1 min, the fluorescence of MaLionG and mitoMaLionR were dropped immediately and then returned to the basal level before heating (Fig. 6B-D). Because the fluorescence of ATP sensors is temperature dependent by itself, the drop of fluorescence during heating was not considered to linked to the only decrease in ATP concentration directly^36^. In the contrast, the delayed recovery in mitochondria was observed even after the disappearance of temperature increment at least, which was reproducible and debatable phenomena (Fig. 6F). Interestingly, mitochondrial ATP took longer time for the recovery to the basal level compared to cytoplasmic one. Also, such differences between them could be identified nearby the heat spot especially (Fig. 6D). The greater the temperature increment, the longer it appears to take for its recovery (Fig. 6E). A mechanistic determinant is assumed that mild thermal effect can perturb the activity of ETC in mitochondria and then it could be recovered reversibly unless it does not exceed the point of the return to induce the cell death^38^. As another aspect, the morphology of mitochondria as well as its location were altered nearby the heat spot, implying that the mitochondria also experienced mechanical stress which in turn might influence the function forcibly (Fig. 6C)^39^. In addition, the quick recovery of cytoplasmic ATP might be reasoned that proteins to play a role for glycolysis are freely moving relatively to compensate the depletion of ATP though the thermal effect on the glycolytic ATP synthesis is still poorly understood.

**Figure 6.**
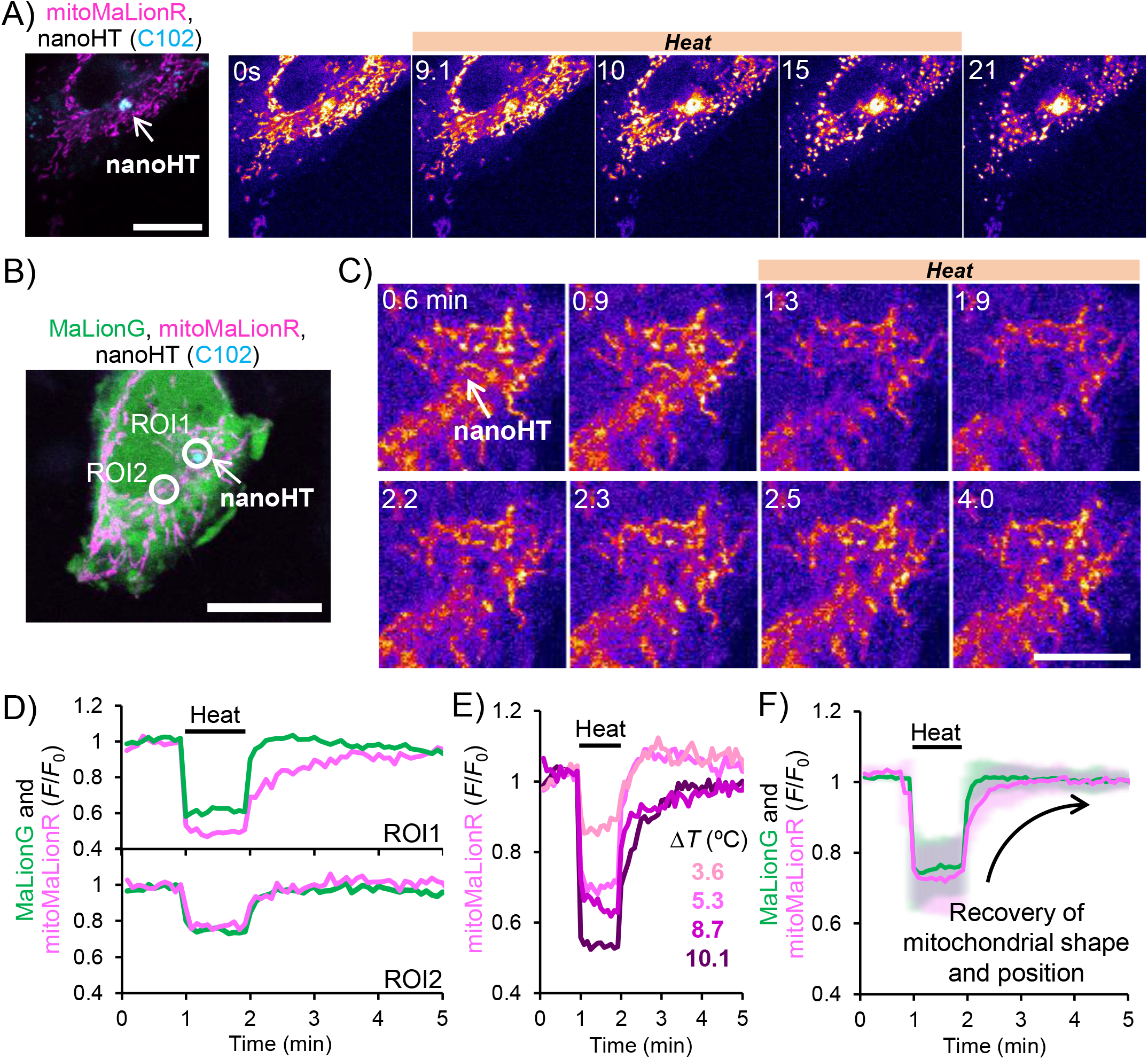
The evaluation of intracellular ATP dynamics during local heating. A) The morphological change of mitochondria occurs after heating at the temperature above the threshold to induce the cell death. Scale bar: 20 μm. B) Fluorescence image of HeLa cell expressing MaLionG (cytoplasmic ATP) and mitoMaLionR (mitochondrial ATP) with **nanoHT** (C102). Scale bar: 20 μm. C) The time course of mitoMaLionR nearby **nanoHT** (the local area of the cell shown in B)). D) The ATP dynamics in cytoplasm (MaLionG) and mitochondria (mitoMaLionR) at ROI1 and 2 of the same cell in B). The heating is 1 min. E) The analysis of mitochondrial ATP dynamics nearby the heat spot like ROI1 in B) in 4 cells at different temperature increments (3.6 to 10.1 °C below the threshold of the cell death). F) The thick lines of MaLionG and mitoMaLionR represent the average of 7 cells with SD at different temperatures.

### Induction of muscle contraction in C2C12 myotube

Photothermal heating strategy has garnered attentions on a means not only to induce the cell death of cancer cells but to other applications. For instance, Oyama et al reported the muscle contraction could be induced by heat, based on the principle that the partial dissociation of tropomyosin with F-actin was promoted thermodynamically^40^. Based on this finding, Marino et al published a success of the remote manipulation of the skeletal muscle contraction using the photothermal effect by the gold-shell nanoparticle^41^. Here, **nanoHT** was applied for the induction of C2C12 myotube contraction at subcellular level. After uptake of **nanoHT** into C2C12 myotube, its cytoplasm was stained with cell tracker green to capture the motion of myotube (Fig. 7A). Repeating on/off of the shutter of an 808 nm laser every 5 sec created the sequential temperature increment, inducing the reversible contraction of the myotube (Fig. 7B and Supplementary movie S1) (ΔT = 10.5±1.4 °C). More importantly, the displacement of the cell involved in the muscle contraction occurred in the limited area of the cell, not whole cell area as has been demonstrated previously^40,41^, which is supported by the kymograph at line A (on the heat spot) and B in the same cell (Fig. 7C-D). Since the distance between line A and B was around 8 μm, the results were also consistent with the analysis of the thermal effect at a few μm scale provided by **nanoHT** (Fig. 4C). The degree of the displacement nearby the heat spot was analyzed from the comparison of the x-z profile before and after heating at the different laser powers (Fig. 7E). The plot of displacement against temperature exhibited the temperature dependency of the degree of the displacement (right panel; Fig. 7E). Upon addition of blebbistatin as an inhibitor for the myosin, the muscle contraction was scarcely observed, suggesting that the contraction was firmly induced due to the alteration of the interaction between tropomyosin and F-actin thermodynamically, not heat-induced physical expansion of cellular volume. The approximate curve correction in the plot seems to represent the exponential feature. Possibly, it might be inferred that the mechanism following the Arrhenius equation was underlying behind the heat induced muscle contraction event though the quantitative analysis is difficult^42^. Through this attempt, it is worth noting that the local heating at subcellular scale enables to manipulate cellular activities at the limited area of a single cell, that has rarely been done using previous technologies.

**Figure 7.**
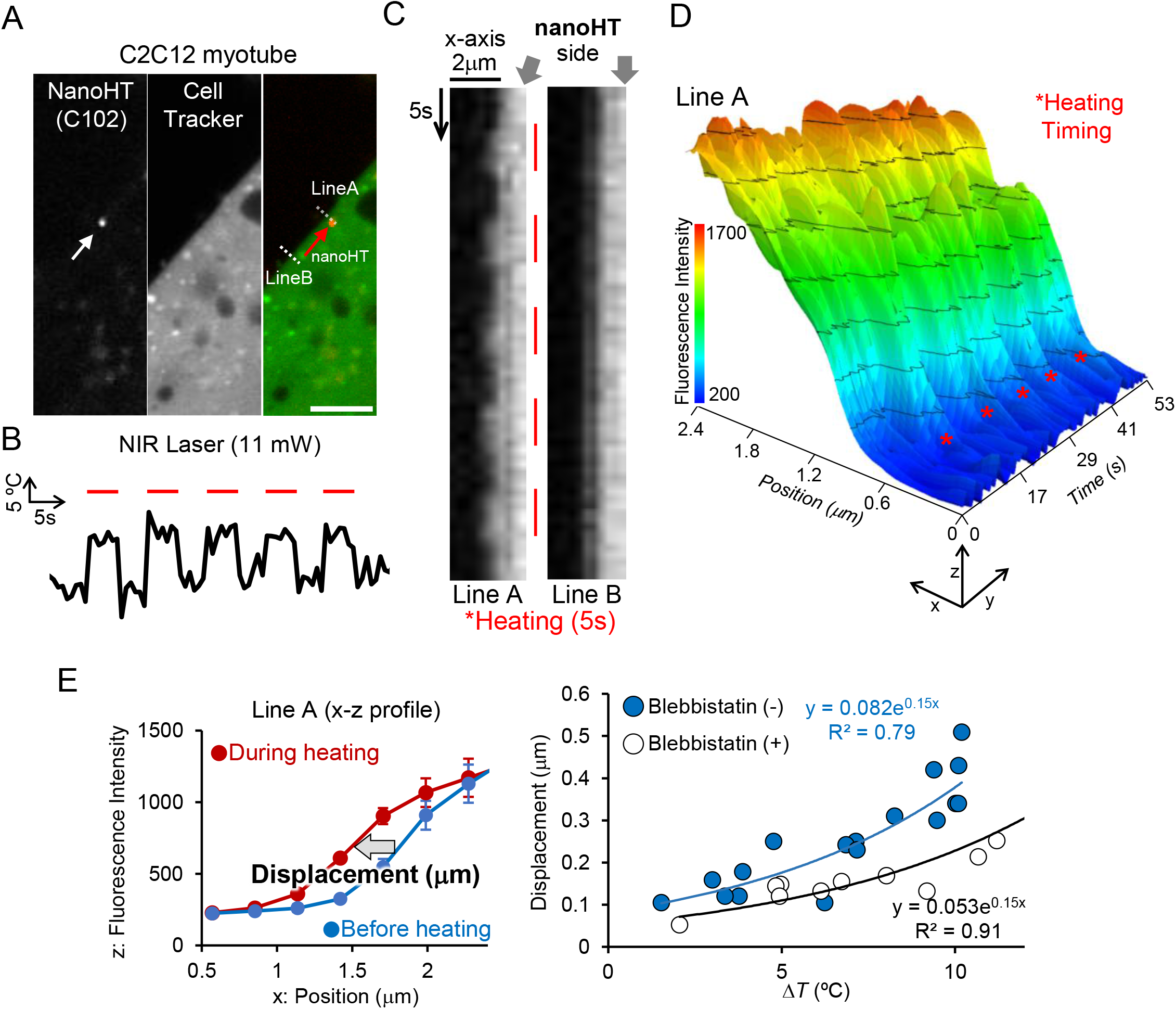
C2C12 myotube contraction induced by sequential heating by **nanoHT**. A) Images of C2C12 myotube with **nanoHT** (red) and Cell Tracker (green: cytoplasm). Scale bar: 10 μm. B) Temperature increments provided by sequential NIR stimulation every 5 sec. C) Kymographs of line A and B as shown in A). D) The dynamic profile of the line A in response to the NIR stimulation. E) Quantitative analysis of the displacement induced by the heating using **nanoHT**. Left panel: the x-z profile of line A. Each dot showed the average with SD during 5 sec. Right panel: the maximum displacement at the x-axis was plotted against varying temperature. The solid line represents the exponential fit.

## Discussion

As one of key things in this paper, we argue more regarding the spatiotemporal dynamics of the temperature distribution created by illuminationg a **nanoHT** with a NIR laser light. Here, the actual spatiotemporal dynamics of heat propagation with non-equilibrium process could not be discussed due to the inherent limitation of a conventional fluorescence imaging^43^. On the other hand, very recent study described that the thermal conductivity inside a complex environment inside a cell (0.11 Wm^−1^K^−1^) is smaller than that of water (0.61 Wm^−1^K^−1^)^44^. If the ease of heat dissipation is governed by the surrounding medium such as water and cytoplasm, the lethal temperature increment of **nanoHT** to reach the steady state would also reflect on the thermal conductivity though the nonequilibrium process could not be captured. However, as a result, the effective differences between in the dish and inside a cell were rarely found regarding the slopes obtained from the correlation between the temperature increment and different laser powers (Fig. 2G and Fig. 3C). That might be due to the limitation of the accuracy and sensitivity of fluorescence thermometry^45,46^. In future, the development of other methodology is required for that purpose, which is beyond fluorescence thermometry^43,47^. While the physical property of thermal elements inside a cell could not be concluded, we showed the distinct difference of the history of temperature distribution during a few seconds of heating. In other words, **nanoHT** is likely to move freely due to the Brownian motion and stir the cytoplasm, which means that even nano-sized heater could provide the thermal effect at submicron scale spatially (Fig. 4C). Therefore, it could be imagined that **nanoHT** collides with some organelles or cytoskeletons, leading to the alteration of cellular activities from subcellular level.

To date, photothermal therapy (PTT) based on NIR modulated nanomaterials to release heat have attracted considerable interests^48^. Nevertheless, the visualization of cellular events with concurrent thermometry in real-time has been scarcely conducted so far. We also stressed the advantage that **nanoHT** is friendly with the other fluorescent indicators, to give the live-cell imaging for cellular activities with the thermometry concurrently. This is practical and powerful enough to contribute to basic research in thermal biology and the development of biomaterials. In most cases, cancer cells undergo apoptosis or necrosis under the elevated temperature around 39-45 °C during hours^49^. Unlikely, we demonstrated subcellular sized heat spot is sufficient to induce the rapid cell death during a few seconds though it is relatively high temperature of around 48°C (Δ*T*=11.4 °C). Yet, the mechanism of the cell death by **nanoHT** still remains under the debate. In particular, the reason why the apoptotic like cell death with the elevation of Apopxin Green was triggered from the local spot and completed during 10 min, which is unlikely to be an intrinsic apoptosis occurring in hours^50^. We also ascertained that this cell death caused by intracellular heating was not characterized by necrosis. This is because determinants to identify necrosis through imaging studies would be totally different from in apoptotic cell death. For example, when the bunch of **nanoHT** was placed on the surface of plasma membrane, we found that the necrotic cell death occurred as proven by the immediate staining of nucleus with PI and scarce fluorescence elevation of Apopxin Green (Supplementary Figure S8). On the other hand, the rapid staining with PI was not detected in the case of intracellular heating by **nanoHT**. Yet, we still could not rule out a possibility that the tiny rapture of the plasma membrane took place from the interior of cell, at the scale of which could not be addressed by an optical microscope. Actually, the damaged cellular membrane resulting from the mechanical stress caused the calcium influx from outside and subsequently annexin V gathers around the inner leaflet at the local spot, which likely matches imaging results (Fig. 5A and 5E)^51^. Since various types of cell deaths have been reported, further studies to unveil the exact mechanism will be required from the viewpoint of cell biology^50,52^.

## Conclusion

In this paper, using **nanoHT**, we could generate subcellular-sized heat spots with the different patterns with varying the amplitude of the laser power and the interval of an 808 nm laser. We successfully demonstrated local heating in not only the induction of rapid cell death but also the manipulation of muscle contraction. From viewpoint of an effective PTT for cancer therapy, a heat-induced cell death for short time is preferred because long-term heating transforms cancer cells to the thermoresistant and ineffective ones for PTT. The heat-induced muscle contraction as the other part is more likely to show great potentials that our concept can extend to various applications. This is because it is based on the thermodynamic alteration of the protein-protein interactions by heating, which can be versatile working principle as a manipulation tool. In future perspective, we do believe that the targeting of **nanoHT** to the desirable places possesses enormous opportunities to alter cellular activities from viewpoint of thermodynamic law, termed “thermodynamic cell engineering”.

## Acknowledgments

This research was supported by Japan Agency for Medical Research and Development (AMED) PRIME (JP18gm5810001), JST FOREST Program (Grant Number JPMJFR201E, Japan), and JSPS KAKENHI (JP20H04702 and JP19H02750).

## Author contributions

SA and F designed this study. SA, F, MS, YH, SI, SS and TK wrote the manuscript. SA and F conducted experiments and analyzed data. MS contributed to the design of microscopic system. All authors discussed the results and commented on the paper.

## Additional information

Supplementary Information including movies accompanies this paper.

## Material and methods

### Materials

Poly(methyl methacrylate-*co*-methacrylic acid)(PMMA-MA)(Mw: 34,000), Vanadyl 2,11,20,29-tetra-tert-butyl-2,3-naphthalocyanine (V-Nc), and Coumarin 102 (C102) were purchased from Sigma-Aldrich. Eu-tris(dinaphthoylmethane)-bis-trioctylphosphine oxide (EuDT) was synthesized according to the previous literature^1^. EBFP-C1 was a gift from Michael Davidson (Addgene plasmid 54738) and B-GECO was also obtained from Addgene. MaLionG and mitoMaLionR were generated in the author’s group (T. Kitaguchi)^2^.

### Preparation and characterization of nanoHT

**nanoHT** was prepared according to the nano-precipitation method^3^. PMMA-MA (5 mg), EuDT (5 mg), V-Nc (0.88 mg) and coumarin 102 (0.25 mg) were dissolved in tetrahydrofuran (THF, 1 mL). 8 mL deionized water was then rapidly added into the organic solution. The mixture was then mixed by gently shaking the bottle. Afterwards, the mixture was left in a fume hood with the bottle uncapped overnight to evaporate tetrahydrofuran. The hydrodynamic diameter of the fabricated nanoparticle was measured by using a Zetasizer ZSP (Malvern). The luminescence properties of the particle were recorded by utilizing a fluorescence spectrophotometer (Cary Eclipse Fluorescence Spectrophotometer, Agilent Technologies) while monitoring the sample temperature with a thermocouple (TES-1310 type-K, TES Electrical Electronic Corp.). The Transmission Electron Microscopy (TEM) image was obtained using Philips CM200 operating at an accelerating voltage of 200 keV. The UV-visible spectroscopy was performed by a UV-vis spectrophotometer (Cary 60 UV-Vis, Agilent Technologies).

### Evaluation of ROS production

H_2_DCF was obtained by deacetylating H_2_DCF-DA for the *in vitro* ROS scavenging assay following reported procedures^4,5^. In brief, H_2_DCF-DA (0.5 mL, 1.0 mM) in methanol was mixed with NaOH (2.0 mL, 0.01 M). The solution was then incubated at 37 °C for 30 minutes with gentle shaking to deacetylate H_2_DCF-DA into H_2_DCF. The mixture was then neutralized with NaH_2_PO_4_ (750 μL, 25 mM) buffer and NaOH (1 mL, 1N) while pH is measured using a pH probe (Sartorious). The non-fluorescence H_2_DCF (117.6 μM) was then stored in the dark at –20 °C. All fluorescence measurements were performed in triplicates. For the *in vitro* assay, a solution of **nanoHT** or AuNR was mixed with H_2_DCF (50 μL, 117.6 μM) and water to achieve the desired final concentration where the temperature increment is similar (0.15 mg/mL of **nanoHT** or 0.03 mg/mL of AuNR) prepared on a 96-well plate. The DCF fluorescence of the sample before illumination (t = 0) was then measured using a by a microplate reader (Infinite M200, Tecan, the excitation and emission wavelengths are 495 nm and 525 nm, respectively). The samples were then exposed to an 808 nm laser at 600 mW for 1 and 3 minutes. Following the illumination, the fluorescence of DCF was then measured again and the relative fluorescence intensity change was then calculated against the intensity before illumination (t = 0). Under conditions in which AuNR and **nanoHT** reached temperature increment, the generation of reactive oxygen species was evaluated.

### Cell viability test

HeLa cells were seeded into a 96-well plate at a density of 5000 cells per well and cultured for 48 hours in culture medium at 37°C under 5% CO_2_ environment. After removal of the culture medium, 10 μL of **nanoHT** (1.2 mg/ml) of different dilution factors (1×, 10×, and 100×) or 10 μL of deionized water (served as blank) and 90 μL of culture medium were introduced to each well and incubated for different time periods (4 hours, 24 hours, and 48 hours). Once the incubation process had completed, 10 μL of MTT (Biotium) solution was added to each well and mixed by gently tapping the plate. The plate was then incubated further for 2 hrs. Afterwards, 200 μL of dimethyl sulfoxide was then added to each well and mixed well until all the formazan salt dissolved. The signal was measured by a microplate reader (Infinite M200, Tecan) and calculated by taking the difference of the absorbance at 570 nm and the background absorbance at 630 nm.

### Photothermal conversion efficiency of nanoHT

The photothermal performance of **nanoHT** was evaluated according to previous literatures^6^. The aqueous solution of **nanoHT** (1 mL) in a quartz cell was illuminated using an 808 nm near-infrared laser for 600 s. A control experiment was carried out similarly using water. A thermocouple (TES 1310 Type-K) was employed to monitor the change in the temperature of the solution every 30 s. An optical power meter (Thorlabs Inc.) was used to adjust the laser output power to 1.0 W·cm^−2^.

The photothermal conversion efficiency (*η*) was determined using reported method as defined in equation (1):

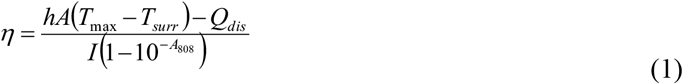

where *h* represents the heat transfer coefficient, *A* is the surface area of the quartz sample cell, *T_max_* is the maximum temperature achieved by laser irradiation, *T_surr_* is the ambient temperature of the environment (23.5°C), *Q_dis_* is the heat dissipation from the light absorbed by the solvent and the quartz sample cell, *I* is the incident laser power (1.0 W·cm^−2^), and *A*_808_ is the absorbance of the sample at 808 nm (0.0675). The value of *hA* was calculated following equation (2):

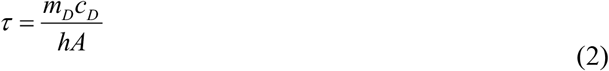

where *m_D_* and *c_D_* are respectively the mass (1.0 g) and heat capacity (4.2 J/g) of the deionized water used to dissolve **nanoHT**. *τ* is the time constant of the sample system. The value of *τ* can be derived following equation (3):

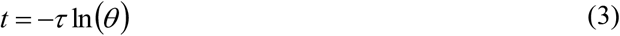

where *t* is the time elapsed after the laser illumination ceases and *θ* is a dimensionless driving force temperature, defined in equation (4) as:

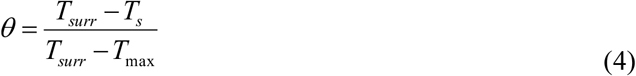

where *T_s_* is the temperature of the sample at a given time *t*.

*Q_dis_* or the heat dissipation from the light absorbed by the solvent and the quartz sample cell can be quantitatively measured following equation (5):

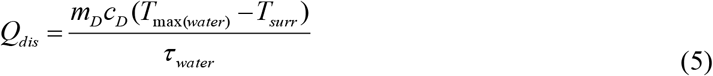

*T_max (water)_* was 25.2°C and *τ_water_* was 430.76 s, thus *Q_dis_* was determined to be 16.6 mW. *T_max_* was 30.6°C and *τ* was 447.88 s, thus *hA* was calculated to be 9.37 mW. Thus, the photothermal conversion efficiency of **nanoHT** (*η*) was determined to be 35%.

### Fluorescence imaging of nanoHT with a NIR stimulation

Fluorescence imaging was performed with an Olympus IX 83 inverted microscope equipped with a FV12-FD camera (Olympus) and an oil immersion objective lens (PLAPON 60×, NA = 1.42). The FV10-ASW 4.2 software (Olympus) was used for controlling camera, filters, and recording data. For a dual color imaging of **nanoHT**, DM405/473 and SDM473 were used as dichroic mirrors and BA430-455 and BA575-675 as emission filters, respectively. For a tricolor imaging of **nanoHT**, DM405/473 and SDM473 and SDM560, BA430-455 and BA490-540 and BA575-675 were used as dichroic mirrors and emission filters respectively (Olympus). For photothermal stimulation during microscopic observation, an Infrared Laser-Evoked Gene Operator (IR-LEGO-100/mini/E) system was introduced to the microscopic setup to allow laser stimulation at 808 nm or 980 nm wavelength (100 mW). In the experiments using a 980 nm laser, iron oxide (Fe2O3) magnetic solution (5 μL) was let dry on a glass-based dish to be used as an external heat source. To obtain the calibration curve of **nanoHT** (normalized ratio of EuDT to C102 vs. temperature), the temperature in the medium was varied from 35 to 48 °C using the microscope temperature-controlled chamber (TOKAI-HIT).

### Cell Culture

HeLa (ATCC^®^ CCL-2^™^) cells were cultured on glass-based dishes in Dulbecco’s modified Eagle’s medium (DMEM) supplemented with fetal bovine serum (FBS, 10%) and penicillin-streptomycin (1%). The cells were grown and kept at 37°C under 5% CO_2_ environment. C2C12 (ATCC^®^ CRL-1772^™^) myoblasts were cultured on collagen I bovine protein (50 μg/mL) coated glass-based dishes in DMEM containing FBS (10%) and penicillin-streptomycin (1%). After the myoblasts reached 100% confluence, they were induced to differentiate into myotubes by replacing the culture media with DMEM supplemented with horse serum (2%) and penicillin-streptomycin (1%). The differentiation process was performed for 7-8 days, with media replacement every 2 days, until the myotubes were developed.

### Temperature mapping of HeLa cells and in the dish

For temperature mapping in the dish, the purified protein of EBFP was used. The purification procedure was followed by the previous paper^7^. The stock solution of EBFP was added to the Hanks’ Balanced Salt Solution (HBSS) buffer so that the effective fluorescence could be observed (the final concentration was adjusted to be 0.1-0.5 mg/ml). For the temperature mapping of cytoplasm in HeLa, HeLa cells (80% confluent) on a 3.5 cm glass-based dish were transfected with 1.0 μg of EBFP (plasmid DNA) using 3 μL of FuGENE HD Transfection Reagent (Promega) in 10 μL of Opti-MEM (Life Technologies Corporation). After transfection, they were kept at 37 °C under 5% CO_2_ for 8 hrs, replaced with a fresh DMEM with 10% FBS, and then incubated at 37 °C for 2 days. The cells were then incubated overnight with 10 μL of **nanoHT** solution added into the medium one day before the observation. The heating experiments during microscopic observation was done using IR-LEGO system (an 808 nm laser) as mentioned above. The trajectory of **nanoHT** in the dish and HeLa was analyzed using the ImageJ software (TrackMate).

### Imaging experiments on the heat-induced cell death (HeLa)

Similar with the experiments regarding temperature mapping of EBFP expressed HeLa cells, **nanoHT** was delivered to HeLa cells through the overnight incubation. For intracellular Ca^2+^ imaging, B-GECO, MaLionG and mitoMaLionR were transfected to HeLa cells instead of EBFP. Prior to imaging experiments to test the cell death, the medium was replaced with 200 μL DMEM and then incubated with a mixture of 2 μL Apopxin Green (Abcam) and 0.4 μL Propidium Iodide (15 mM, Thermo Fisher) for 15 minutes at 37 °C under 5% CO_2_. The heating experiments were done using IR-LEGO system (an 808 nm laser). To evaluate whether it induce the cell death or not, the laser power was varied from 8.8 to 11.2 mW.

### Imaging experiments on heat-induced muscle contraction (C2C12)

Prior to imaging, the myotubes were incubated overnight with 10 μL of the **nanoHT** solution added into the medium one day before the observation. The cells were stained with 100 nM Calcein, AM (Thermo Fisher) to visualize the myotubes upon 808 nm laser illumination by IR-LEGO. To inhibit myotube contraction, a myosin inhibitor, Blebbistatin (Thermo Fisher, 25 μM) was introduced into the medium. The kymograph was obtained from the analysis with the ImageJ software. The 3D image of the muscle contraction was generated from the RINEARN Graph 3D Software.

