## Supplemental Figures for "Photothermal Dye-based Subcellular-sized Heat Spot Enabling the Modulation of Local Cellular Activities"

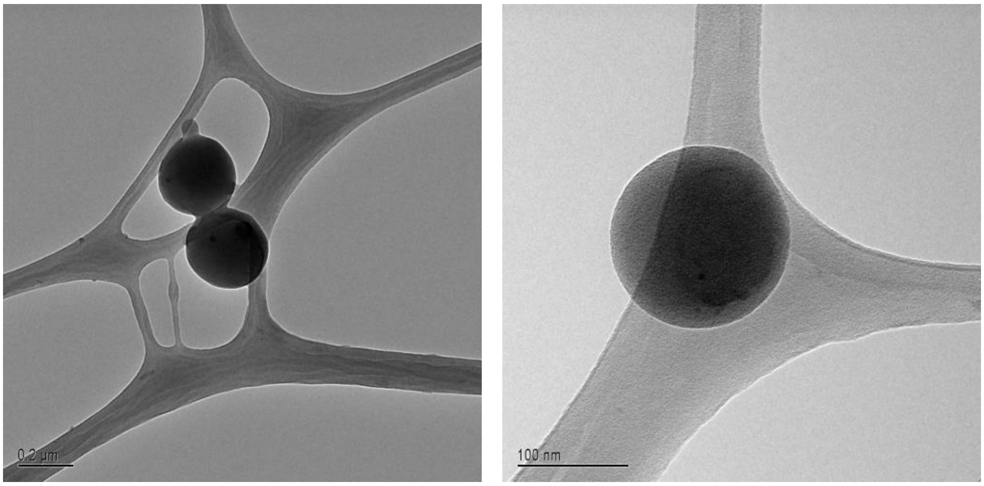


**Figure S1**. TEM images of **nanoHT**.

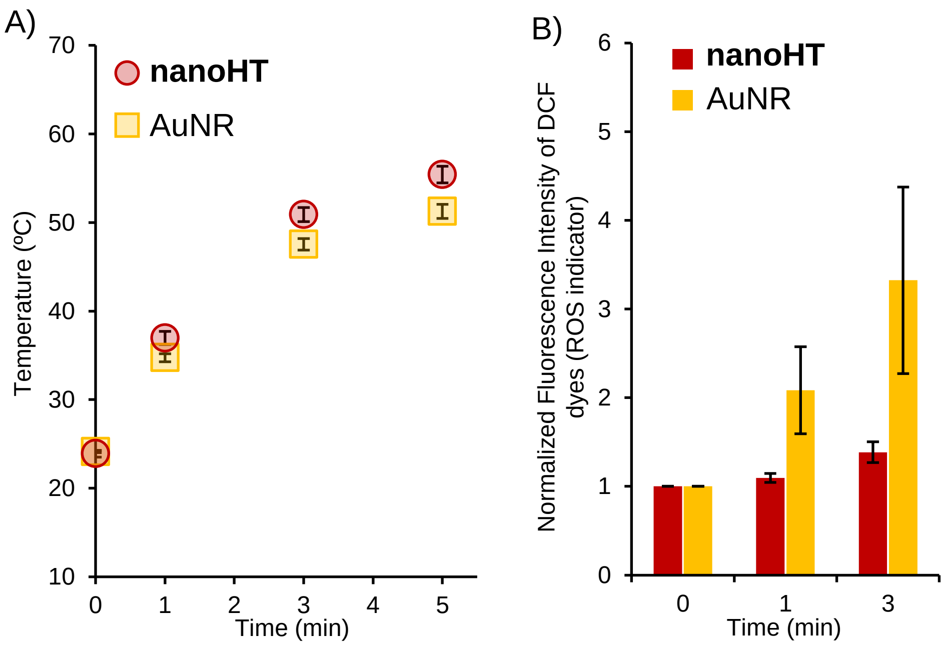


**Figure S2.** Evaluation of reactive oxygen species (ROS) production in comparison with **nanoHT** and AuNR (commercially available gold nanorods: 10 nm x 41 nm, Sigma Aldrich #716820). A) The aqueous solution of **nanoHT** (0.15 mg/ml) and AuNR (0.03 mg/ml) showed almost same temperature increment respectively. The irradiation laser power was 600 mW (808 nm). B) Under the same condition with A), the ROS production was evaluated using a DCF dye. Each data point represents the average of three independent experiments. Error bars, SD (n=3).

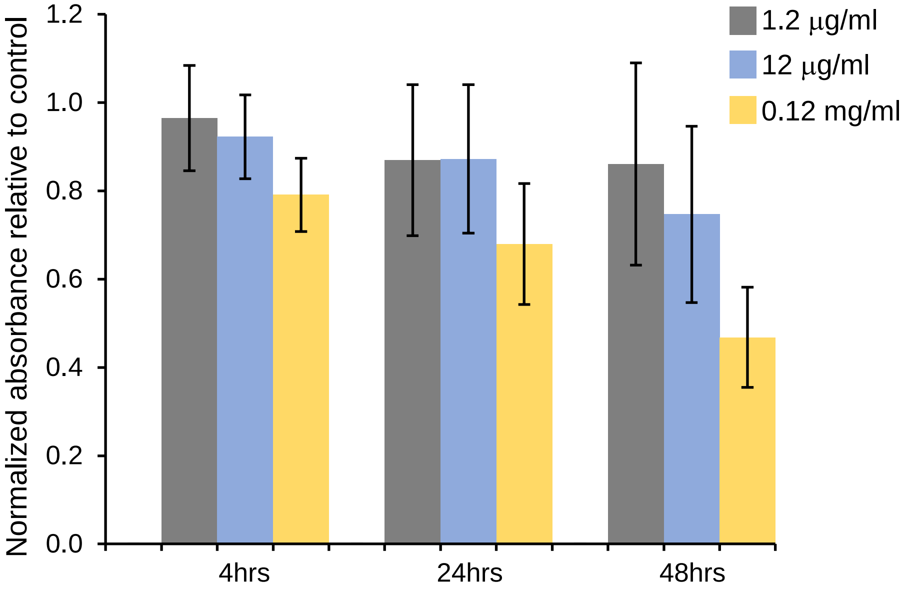


**Figure S3.** Cell viability test of **nanoHT**. MTT assay was done with three different concentration of **nanoHT** (1.2 μg/ml, 12 μg/ml, 0.12 mg/ml).

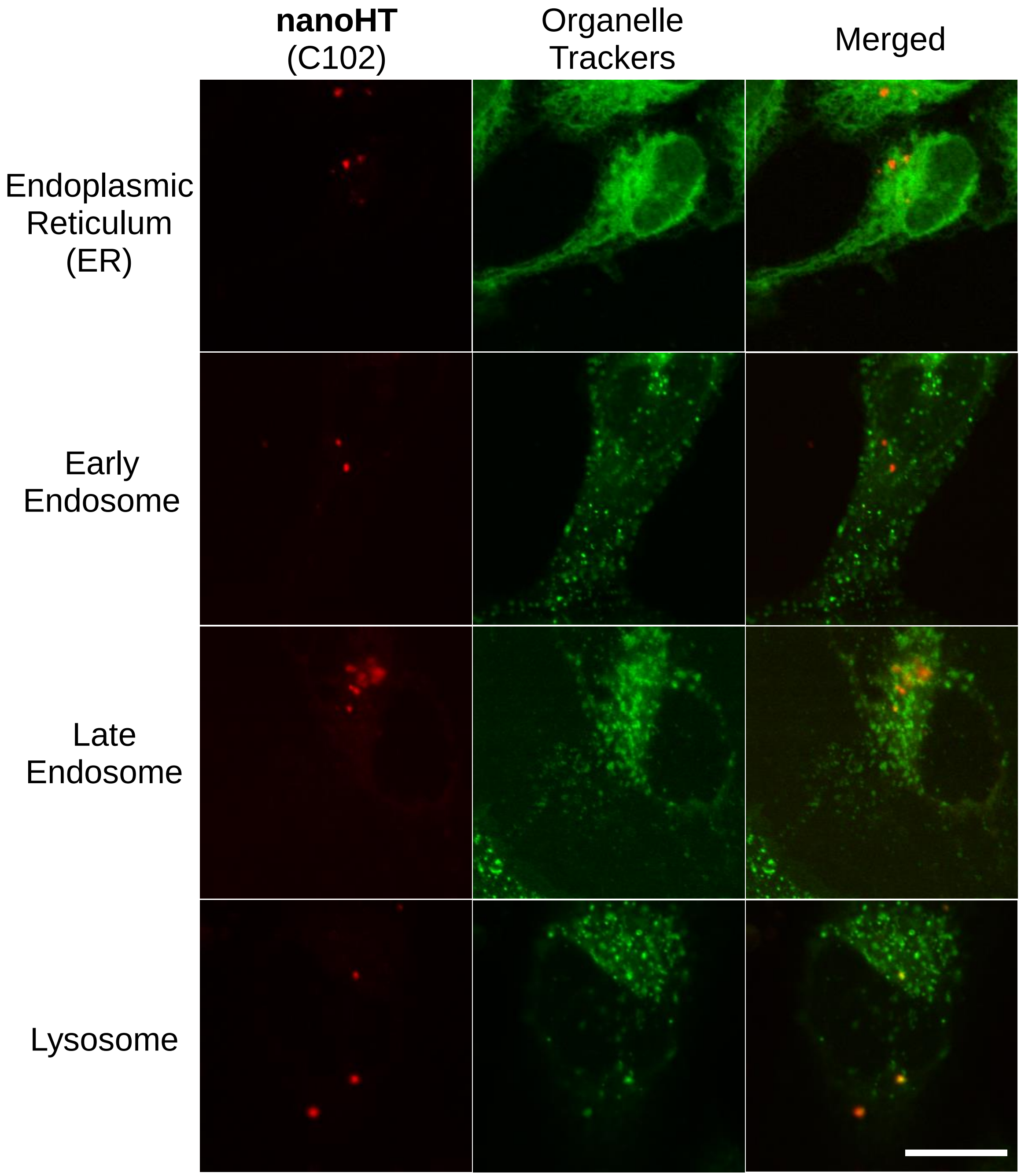


**Figure S4**. Colocalization tests of **nanoHT.** Scale bar: 20 μm

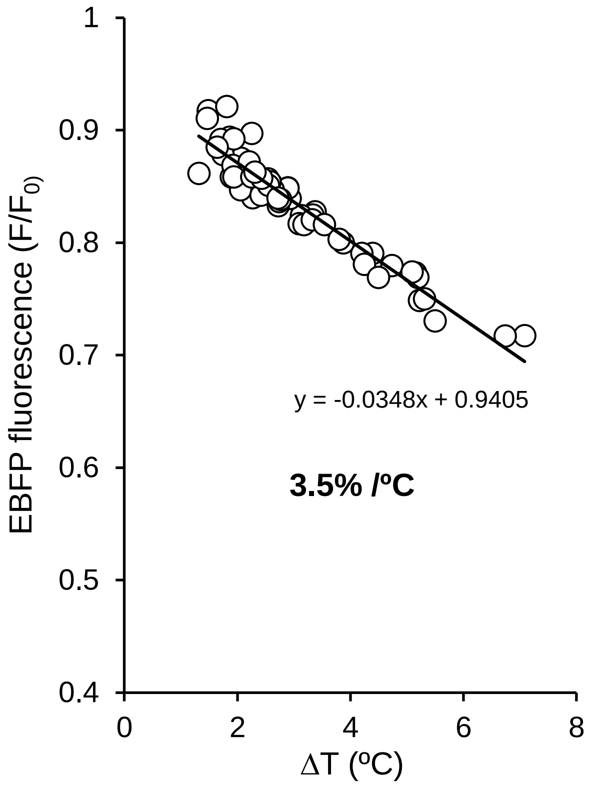


**Figure S5**. The calibration curve of the purified EBFP solution (EBFP-C1).

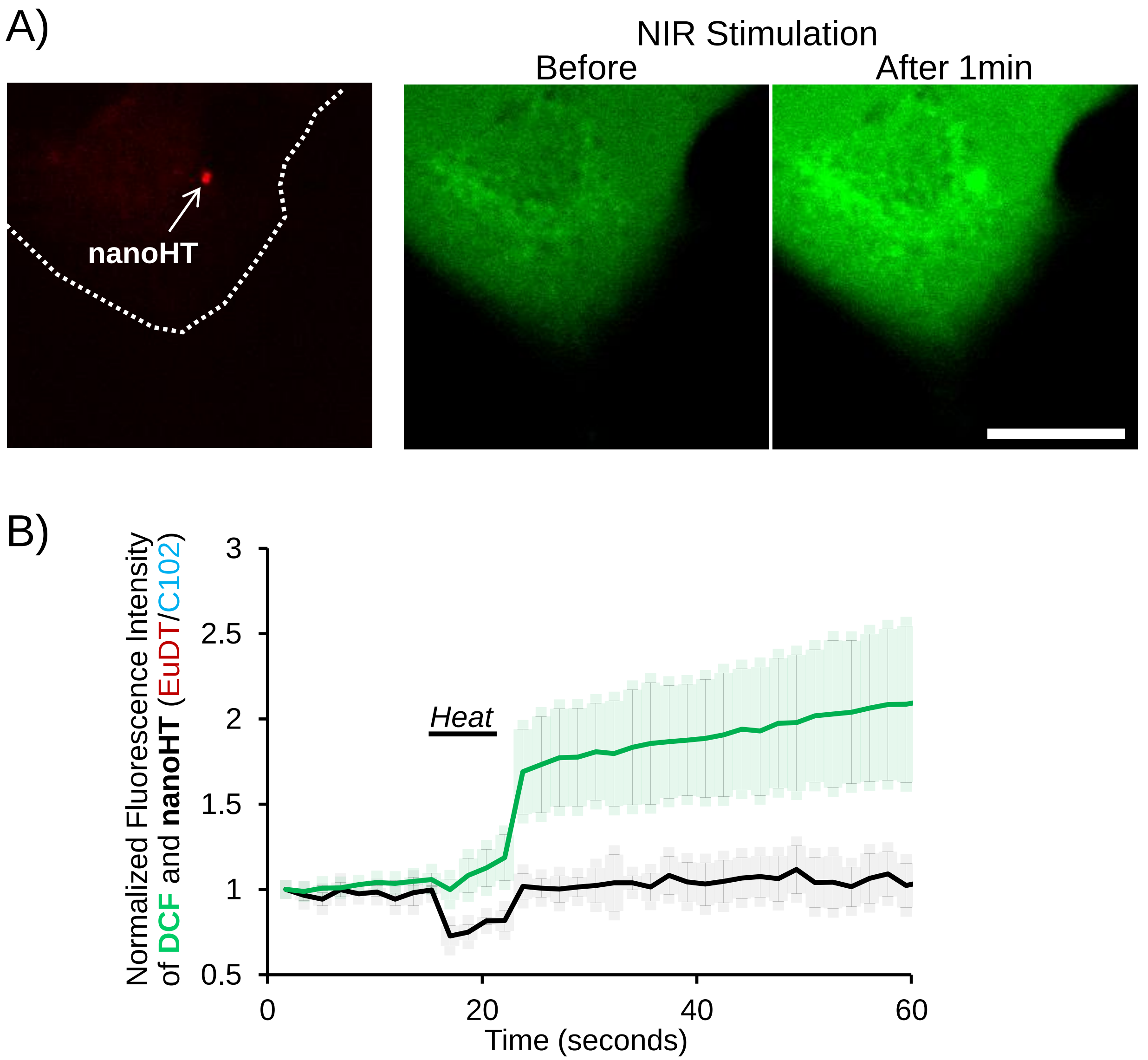


**Figure S6** Intracellular ROS Imaging using a DCF dye. A) Images of location of **nanoHT** and DCF before and after a NIR stimulation (0 and 60 sec.). Scale bar: 20 μm. B) The time course of fluorescence of DCF and **nanoHT** (the average of representative 3 cells in the independent dishes with SD). The average of temperature increment was 12.7 ± 0.4 ºC, which was higher than the threshold inducing the cell death.

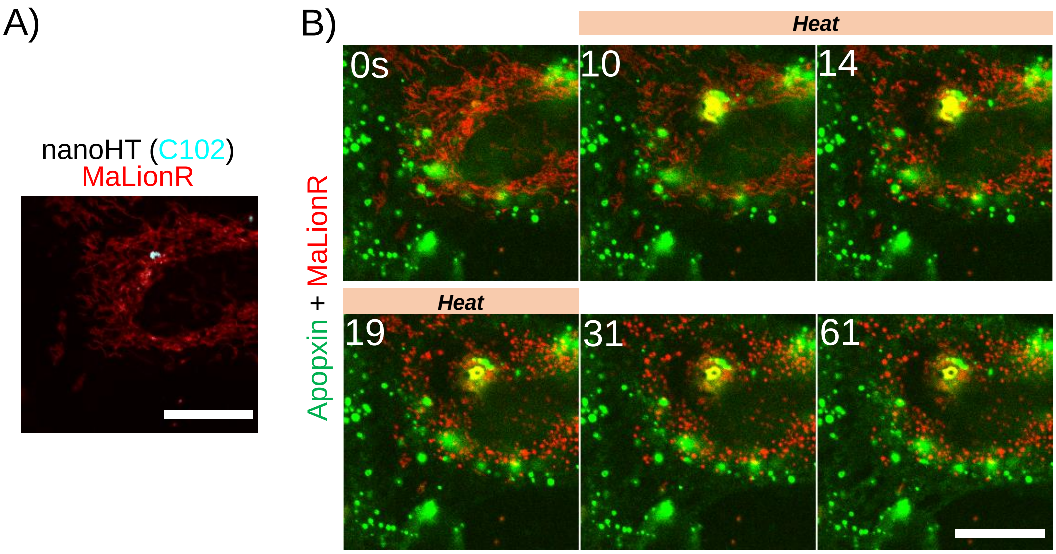


**Figure S7**. Dual imaging of Apopxin (green) and mitochondrial ATP (mitoMaLionR). A) A single **nanoHT** (C102, nanoHT) was observed in the network of mitochondria. B) Dual imaging of Apopxin and MaLionR. Scale bar: 20μm

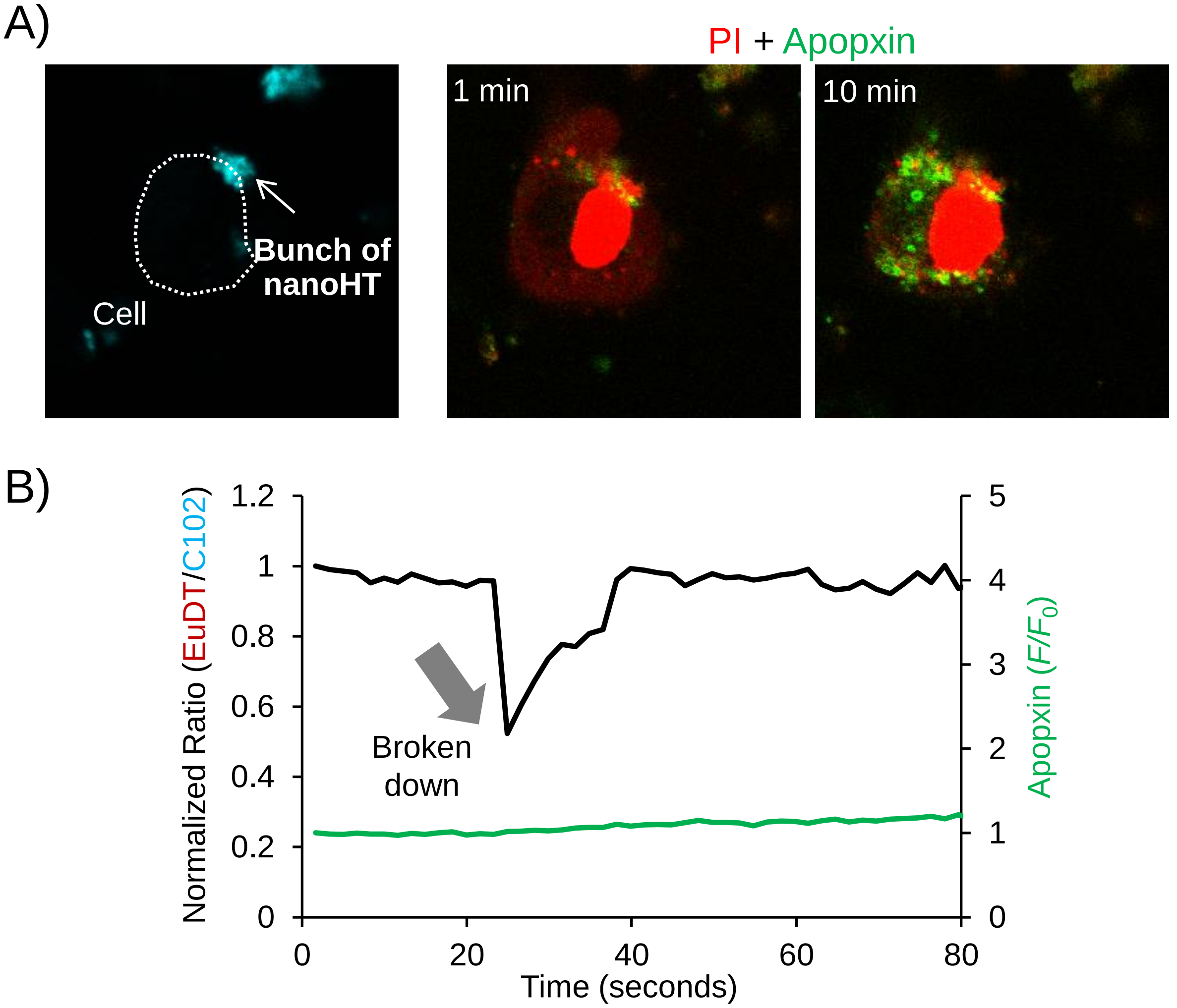


**Figure S8**. Bunch of **nanoHT**s outside induced the necrotic cell death. A) A bunch of **nanoHT**s are located at the periphery of cellular membrane (nanoHT, C102 channel). Cells were stained with PI (red) and Apopxin (green). After the NIR illumination, the fast fluorescence increase of PI was observed while Apopxin was negligible, which was identified to the necrotic cell death. B) The fluorescent thermometry was affected by the broken bunch of **nanoHT**s.
